# Sex Steroid Hormone Signaling Tunes Metabolic and Neuronal Programs in Human Cortical Development

**DOI:** 10.64898/2026.05.16.725519

**Authors:** Hanna E Berk-Rauch, Laura-Yvonne Gherghina, Lilin Huang, Kelsey Hennick, Tomasz J. Nowakowski, Andrea H. Brand, Aravinda Chakravarti

## Abstract

Autism spectrum disorder (ASD) exhibits a profound male biased sex ratio. While numerous genes have been implicated in ASD, the *functional basis* of this sex difference is unclear. One enticing hypothesis is genome-wide transcriptional regulation through estrogens and androgens. While hormone-mediated transcription is well-studied in reproductive tissues, its role in cortical development is poorly defined. Thus, we profiled androgen (AR) and estrogen (ESR1/ESR2) receptor expression in primary human cortex and developmental stage-matched organoid models by single-cell RNA-seq. AR was primarily expressed in radial glia and intermediate progenitors while ESR1/ESR2 was more broadly distributed across multiple cell types of the developing cortex, although with the highest expression in radial glia. To study their genetic effects, we exposed iNeurons and cortical organoids to physiological levels of dihydro-testosterone (DHT) and estradiol (E2). DHT consistently up-regulated oxidative metabolism programs enriched in progenitor cells and down-regulated neuronal maturation pathways, while E2 exhibited a much more attenuated effect. The presence of DHT reduced *NTRK2* (TrkB) expression, correlating with expression in developing cortex where *NTRK2* had significantly higher expression in progenitor cells of the female cortex, which is also reflected in the increased expression of AR in radial glia. Together, these data indicate that in developing human cortical lineages, sex hormones act as selective, cell-state– dependent modulators that tune metabolic and maturation programs rather than broadly reprogramming the genome. Thus, the effects of variation in transcriptional regulation through estrogens and androgens are likely to be minor, but not absent, in ASD.

## Introduction

Many disorders and associated traits exhibit significant sex differences in incidence or severity, some profound, yet the mechanisms underlying these differences remain incompletely understood. The classical explanations of extreme sex-bias are sex-limited expression, such as in the reproductive tissues, or X-linked inheritance but most disorders have a sex-bias not limited to one sex. Thus, alternative non-sex-limited hypotheses need to be considered. Neurodevelopmental disorders (NDDs), including autism spectrum disorder (ASD), belong to this latter category since ASD has a 2:1 – 4:1 higher prevalence in males than females(1),(2). While genes contribute substantially to ASD and other NDDs, only a minority of syndromic cases (e.g., Rett syndrome, Fragile X syndrome) are due to single X-linked mutations. Conversely, hundreds of autosomal risk genes have been implicated(3,4), suggesting that differences in sex chromosome gene dosage alone are unlikely to fully explain observed sex biases. We hypothesize that sex hormones can bias disorder gene expression differentially in males and females and contribute to the sex bias.

These observations motivate a mechanistic understanding of how sex hormones can modulate human cortical development, a major target tissue for ASD(5), in ways that could contribute to sex-biased neurodevelopment and its pathologies, including but not limited to ASD. Sex hormones provide a plausible cis-regulatory mechanism for sex differences by modulating gene expression through their cognate receptors. Estrogens and androgens are established genomic regulators in reproductive tissues via their receptors ESR1, ESR2, and AR, which bind cis-regulatory elements to control target gene transcription(6–9). During and after puberty, hormone-dependent gene regulation is a key determinant of sex differences across diverse physiological systems (e.g., immune and cardiovascular(10,11)). In contrast, hormone actions during prenatal brain development are less clearly defined beyond the mid-gestation testosterone surge in male fetuses associated with sexual differentiation(12). Metabolomic studies suggest that estrogens and androgens are present in prenatally developing tissues(13), and placental androgen levels have been associated with increased ASD risk in some cohorts(12,14,15).

Work in rodent models demonstrates hormone-dependent regulation in specific neuronal populations. For example, androgens regulate dimorphic development of peripheral tissues via transcriptional control of targets such as the BDNF receptor NTRK2(16), and estrogens can act as genome-wide regulators of masculinization in subcortical nuclei (e.g., BNST)(17). In hippocampal neurons, estrogen alters protein expression and synaptic function(18). While these findings are informative mechanistically, important differences in expression and developmental physiology exist between rodents and humans. Beyond well-established differences in X-inactivation(19,20), brain masculinization in mice is largely mediated by aromatization of testosterone to estrogen(17), whereas non-human primate and human cell models support a prominent role for testosterone-derived dihydrotestosterone (DHT) with less clear effects from aromatization(21,22). Therefore, direct studies of human cortical systems are needed for more relevant insight.

ASD has widespread phenotypic effects across the brain, with nearly every brain region vulnerable to cellular and network alterations. However, autism-associated gene expression is enriched in the human cortex during mid-gestation(5), implicating this tissue as a point of convergent genetic effects in development. Furthermore, this period of development has implications for processes underlying ASD, as it coincides with the bulk of neurogenesis. Histological studies in rodents report sex hormone receptor expression in the developing cortex(23,24), with some evidence for ESR1 in progenitor classes and proliferative responses to exogenous estrogen. Evidence for AR, ESR1, and ESR2 expression in human prenatal cortex is limited, though AR expression has been observed in first trimester cortex(25), while ESR1 and ESR2 are suggested to be expressed in the first (ESR1) and second (ESR2) trimester cortex, respectively(26). Sustained androgen exposure can increase proliferating radial glia and excitatory progenitor populations in cortical organoids(22), whereas extended estrogen exposure reduces proliferation and promotes maturation(27), suggesting opposing influences on shared developmental pathways. Acute DHT exposure elicits transcriptional responses in both 2D and 3D neuronal cell models, although the magnitudes and target genes are variable across systems(22,28).

To address these knowledge gaps, in this study we characterize gene expression of AR, ESR1, and ESR2 in the prenatal human cortex and in complementary cortical culture models. Furthermore, we directly test acute transcriptional responses to estrogen and androgen exposure during normal cortical development. We find that sex hormone receptors are present in the primary cortical system and in *in vitro* models. Consistent with restricted receptor availability and developmental context, both hormones elicit targeted transcriptional effects—with androgen exposure in particular down-regulating neuronal maturation genes and up-regulating metabolic and cell-cycle signatures, with targets like NTRK2(16). These results suggest that in developing cortical lineages, sex hormones exert selective, cell-state–dependent transcriptional influences, with androgen effects concentrated in progenitor cells and estrogen promoting maturation, rather than broad genome-wide reprogramming.

## Results

### Sex hormone receptors are expressed in the developing cortex and in culture models

The transcriptional effects of sex hormones are mediated by their canonical nuclear receptors: the androgen receptor (AR) encoded on the X chromosome, and the autosomal estrogen receptors 1 and 2 (ESR1 and ESR2). Because antibody quality and specificity for these proteins are variable²⁹, ³⁰, we assessed receptor expression at the transcript level using complementary single-cell and single-nucleus RNA-seq datasets spanning human cortical development.

### Cell-Type-Specific Expression During Mid-Gestation

We first examined receptor expression during mid-gestation (gestational weeks 16-24), when testosterone rises rapidly in males^31^, using scRNA-seq from primary human cortex tissue (38,526 cells analyzed).(29) Across major cortical cell classes (**Figure 1A**), transcripts for *AR*, *ESR1*, and *ESR2* were detectable, although each receptor was expressed at relatively low levels overall (**Figure 1B**). Detection rates during mid-gestation were: *AR* 1.0%, *ESR1* 4.6%, *ESR2* 15.2% of cells. This sparse expression pattern is consistent with the known biology of nuclear hormone receptors in cortical tissue, which typically operate at low copy numbers but achieve biological effects through catalytic amplification of transcriptional responses.

**Figure 1.**
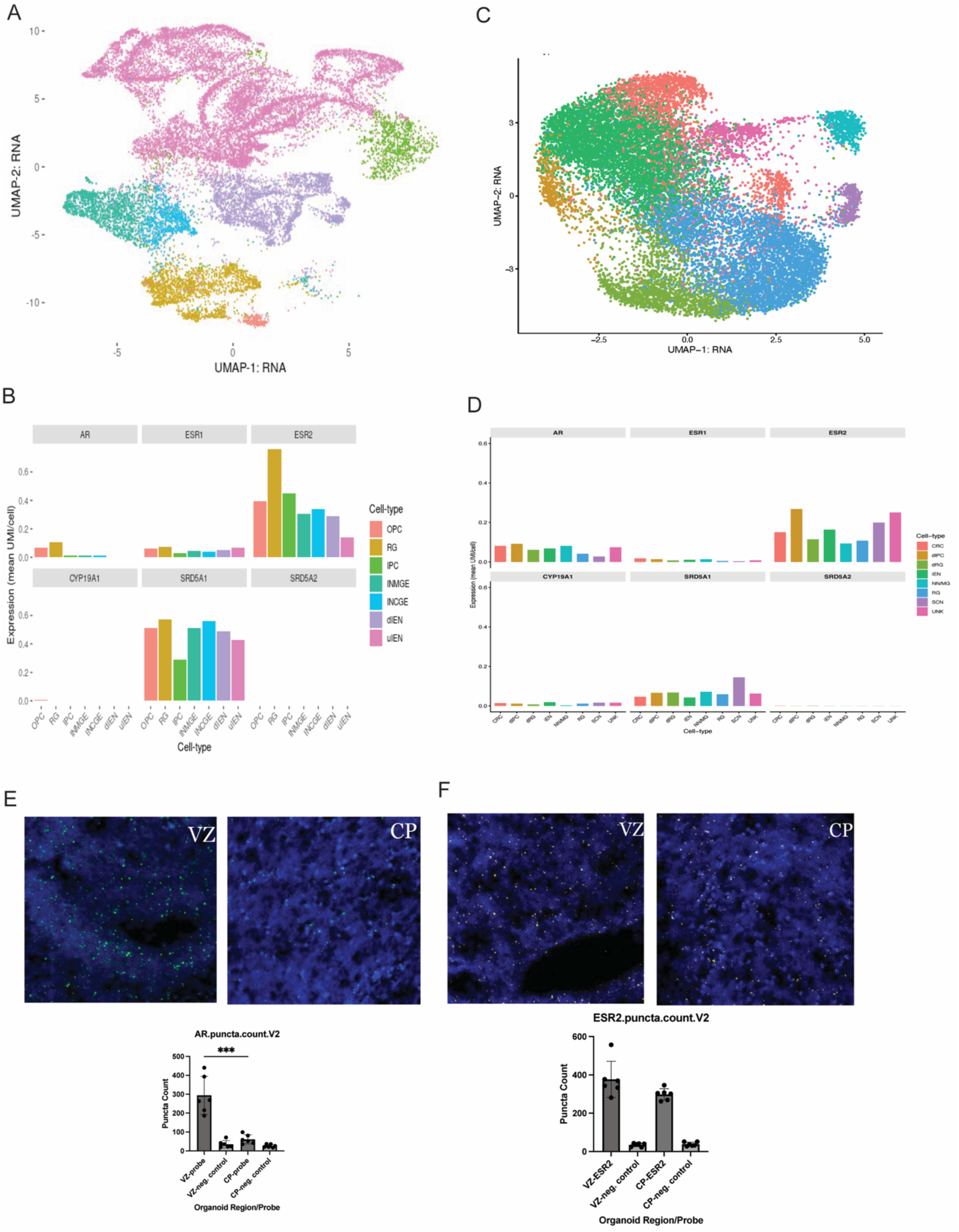
Sex hormone receptors are sparsely expressed and cell-type– specific in prenatal developing cortex and cortical organoids. **(A)** UMAP visualization of 38,526 single cells from developing human cortex (gestational weeks 16-24) colored by cell type. Major cell classes: radial glia (RG, n=2,499), intermediate progenitors (IPC, n=2,009), dividing cells (Div, n=1,203), excitatory neurons upper layer (UL-EN, n=21,094) and deep layer (DL-EN, n=6,170), interneurons (IN, n=4,821), oligodendrocyte precursor cells (OPC, n=469), microglia (MG, n=163), endothelial cells (Endo, n=98). **(B)** Bar plots showing mean normalized expression (UMI/cell) of sex hormone receptors (*AR, ESR1, ESR2*) and steroidogenic enzymes (*CYP19A1, SRD5A1, SRD5A2*) across cortical cell types in developing cortex; color legend at right. **(C)** UMAP of single-cell RNA-seq from DIV20 cortical organoids generated with a semi-guided cortical protocol, colored by inferred cell type. Organoids recapitulate major progenitor and early neuronal populations observed in primary tissue (cajal-retzius cells [CRC], dividing intermediate progenitors [dIPC], dividing radial glia [dRG], immature excitatory neurons [iEN], nonneuronal/microglia [NN/MG], radial glia [RG], subcortical neurons [SCN], unknown [UNK]). **(D)** Bar plots showing mean normalized expression (UMI/cell) of sex hormone receptors (*AR, ESR1, ESR2*) and steroidogenic enzymes (*CYP19A1, SRD5A1, SRD5A2*) across cortical cell types across organoid cell types; color legend at right. **(E)** RNAscope detection of *AR* transcripts (green puncta) in organoid ventricular zone (VZ)-like and cortical plate (CP)-like regions (representative images; DAPI in blue). Bottom: quantification of *AR* puncta per region and probe (mean ± s.e.m.), demonstrating higher *AR* signal in VZ-like zones as compared with CP and negative controls (***P < 0.001, test). **(F)** RNAscope detection of *ESR2* transcripts in VZ and CP of organoids. Bottom: quantification of *ESR2* puncta per region and probe (mean ± s.e.m.), showing robust *ESR2* signal in VZ and CP and low background in control probes.

*AR* expression was preferentially enriched in radial glia (RG; 8.2% positive cells, 14.70-fold enrichment vs other cell types, p = 1.1×10⁻¹³¹, hypergeometric test) compared with other cortical cell types (**Figure 1B**), whereas *ESR1* and *ESR2* were distributed more broadly across multiple neuronal and glial classes, including interneurons (IN; ESR2: 18.1% detection), layer 2-6 excitatory neurons (EN; ESR2 range: 8.8-17.2%), intermediate progenitors (IPC; ESR2: 28.4%), and dividing cells (Div; ESR2: 45.9%) (**Figure 1B**). Notably, ESR1 showed more uniform distribution across cell types (coefficient of variation = 0.25) compared to the cell-type-restricted pattern of AR (CV = 1.02), while ESR2 showed intermediate variability (CV = 0.44) based on UMI/cell expression levels. This distribution pattern is consistent with a model wherein androgen signaling capacity is concentrated within progenitor compartments—where it may influence proliferation, fate specification, or neurogenic timing—while estrogen receptors are more diffusely expressed across cortical lineages, potentially mediating pan-cellular effects on neuronal maturation and circuit assembly.

To assess the local metabolic capacity for sex hormone processing, we additionally examined expression of key steroidogenic enzymes: aromatase (*CYP19A1*), which converts testosterone to estradiol, and 5α-reductases 1 and 2 (*SRD5A1*, *SRD5A2*), which convert testosterone to the more potent androgen dihydrotestosterone (DHT). The vast majority of testosterone in target tissues is locally converted to either estradiol or DHT rather than acting directly³². Aromatase (*CYP19A1*) was largely undetectable (0.2% of cells), *SRD5A1* was widely expressed (26.4% detection), whereas *SRD5A2* was minimal (0.2% detection) (**Figure 1B**). This expression pattern suggests that testosterone exposure in developing cortex is predominantly processed to DHT rather than estradiol, implying that androgenic (AR-mediated) rather than estrogenic (ESR1/ESR2-mediated) signaling may be the primary mode of testosterone action during this critical developmental window.

### Developmental Trajectories from Development Through Adulthood

To examine developmental dynamics across a broader temporal range, we analyzed published single-nucleus RNA-seq (snRNA-seq) data spanning 40 developmental stages from prenatal development (4th gestational month, ∼16 post-conceptional weeks) through late adulthood (54 years) in both male and female specimens(30) (normalization and modeling detailed in Methods; 709,372 cortical nuclei).

*ESR2* demonstrated significantly higher expression than both *ESR1* and *AR* across all developmental stages (ESR2 vs ESR1: mean log-normalized expression 0.132 vs 0.079, V = 731, p = 3.2×10⁻⁶, Cohen’s d = 1.08; ESR2 vs AR: 0.132 vs 0.057, V = 775, p = 3.1×10⁻⁸, Cohen’s d = 1.52; paired Wilcoxon signed-rank test; **Supplemental Figure 1A**). *ESR1* expression was also significantly elevated compared to *AR* (V = 681, p = 1.4×10⁻⁴, Cohen’s d = 0.85). These large effect sizes (Cohen’s d > 0.8) indicate substantial biological differences despite low absolute expression levels. Developmental trajectory analysis revealed distinct temporal patterns for each receptor (**Supplemental Figure 1A**). *ESR2* expression peaked during early prenatal development (4th-6th gestational months; mean = 0.201) and remained elevated relative to other receptors throughout the lifespan. *ESR1* levels were relatively stable across prenatal and postnatal stages (coefficient of variation = 0.32), with modest fluctuations during childhood and adolescence. In contrast, *AR* expression peaked from mid-gestation through the first postnatal year (mean = 0.069) compared to later developmental stages (mean = 0.050), though this difference was marginally significant (t(15.6) = -1.77, p = 0.097, Welch’s t-test). The highest *AR* expression was detected in early postnatal (0.154), newborn (0.102), and 9th gestational month (0.098), paralleling known testosterone dynamics during the late gestational surge and the “mini-puberty” of infancy³⁴.

Despite well-established sex differences in circulating hormone levels, we did not detect significant sex differences in cortical receptor expression at the bulk tissue level after correction for multiple testing (*AR*: median difference = -0.006, V = 287, p = 0.16, FDR > 0.05; *ESR1*: median difference = 0.011, V = 436, p = 0.68, FDR > 0.05; *ESR2*: median difference = 0.015, V = 453, p = 0.58, FDR > 0.05; paired Wilcoxon signed-rank test across 27-30 matched developmental stages; **Supplemental Figure 1B**). Effect sizes were minimal for all three receptors (median differences <0.02 log-normalized expression units, corresponding to <5% linear-scale difference). This lack of sex dimorphism persisted across developmental periods, including the prenatal/early postnatal window when *AR* expression peaked (**Supplemental Figure 1C**). However, low expression levels and cellular heterogeneity inherent to bulk tissue analysis limit statistical power for detecting cell-type-specific sex differences (see Methods for power analysis).

### Cortical Organoids Recapitulate Sex Hormone Receptor Expression Patterns of the Developing Human Cortex

To establish an experimental platform for investigating hormone-responsive transcriptional programs, and to benchmark *in vitro* findings against primary tissue, we profiled 3D cortical organoids generated using a semi-guided cortical differentiation protocol(31). Single-cell RNA-seq performed at DIV20 yielded ∼30,000 cells across three biological replicates, spanning progenitor and early neuronal states as annotated using established cortical lineage markers(32) (Fig. 1C). Expression patterns of *AR*, *ESR1*, and *ESR2* in organoids closely mirrored those observed in primary tissue, both in normalized transcript abundance and in the fraction of receptor-positive cells within specific lineages (Figure 1D). Cell-type specificity was also similar, with receptor enrichment patterns recapitulating those detected *in vivo*. Independent validation using RNAscope confirmed receptor transcript detection and spatial localization within organoid structures (Figures 1E, F).

Collectively, these data demonstrate that cortical organoids preserve key features of sex hormone receptor expression seen in the developing human cortex, supporting the presence of hormonally responsive transcriptional capacity within defined progenitor and neuronal compartments and validating this system for downstream perturbation studies.

### Exposure to sex hormones elicits a restricted transcriptional response in cortical culture models

To identify the acute hormone effects on neuronal transcription, we treated induced male neurons (iNeurons)(33) with estradiol (E2) or dihydrotestosterone (DHT) at 5, 10, 20, and 100 nM for 24 hours, alongside vehicle controls. We chose a 24-hour exposure to maximize detection of direct receptor-mediated transcriptional responses, rather than secondary physiological changes that have been reported under longer treatments(22,27). Additionally, since iNeurons lack aspects of intact tissue physiology, we performed analogous treatments in 3D cortical organoids (semi-guided differentiation(31)). Because sex hormone receptor expression is highest in radial glia and immature neurons (Figure 1), we treated organoids at two early differentiation time points (DIV20 and DIV40) to capture immature and developing cell types. Organoids were exposed also for 24 hours to 10, 20, or 100 nM DHT or E2 with matched vehicles.

Across doses, DHT elicited a stronger transcriptional response than E2, but in both cases the number and effect sizes of differentially expressed genes (DEGs) were modest (median 48 DEGs per experiment (range: 1-254); Figures 2A, C). Dose ranges were aligned with prior neuronal/cell-line studies(22,28) and within the physiological/near-physiological range(34),(35). To compare the transcriptional control of DHT and estrogen, we quantified differentially expressed genes (DEGs, padj<0.05) across seven independent experiments (iNeurons at 5, 10, 20, 100nM and Cortical Organoids at 10, 20, 100nM) for each hormone. DHT treatment resulted in detectable transcriptional responses (≥1 DEG) in 6 of 7 experiments, compared to 4 of 7 for estrogen. Notably, DHT elicited robust responses (≥10 DEGs) in 71% of experiments (5/7), while estrogen produced robust responses in only 29% (2/7). Among responsive experiments, DHT produced a median of 48 DEGs (range: 1-254) while estrogen produced a median of 8 DEGs (Figures 2B, C; range: 1-85). Although this ∼6-fold difference in median response did not reach statistical significance (p = 0.39, Wilcoxon rank-sum test), the higher response rate and greater magnitude of transcriptional changes suggest DHT elicits a more potent and consistent transcriptional response than estrogen in this cellular context (Figure 2A).

**Figure 2.**
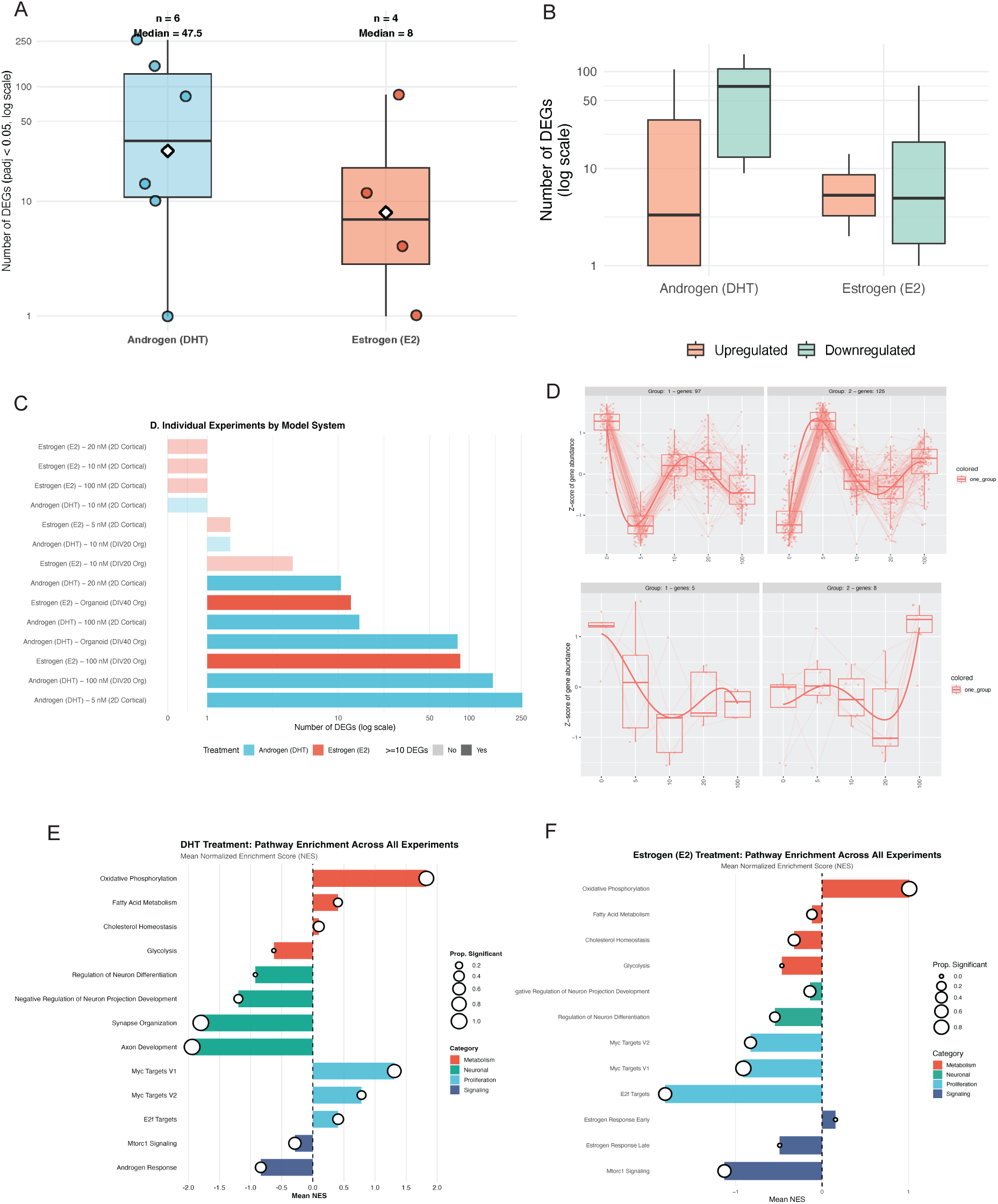
Acute androgen and estrogen exposure produce consistent targeted transcriptional responses across cortical models. **(A)** Number of differentially expressed genes (DEGs; FDR < 0.05) per experiment for 24 h DHT (blue) versus E2 (orange) treatment as compared with vehicle in 2D iNeurons and 3D organoids. Each point is a single RNA-seq experiment; boxplots summarize the distribution. DHT exposures yield more DEGs overall (median ∼47.5) than E2 (median ∼8). **(B)** Numbers of up-regulated (orange) and down-regulated (green) DEGs for DHT and E2 treatments (log_₁₀_ scale). For both hormones, down-regulated genes generally outnumber up-regulated genes, indicating a predominantly repressive acute response. **(C)** Summary of DEG counts in individual experiments stratified by treatment and model system. Horizontal bars (log_₁₀_ scale) show total DEGs for each hormone, dose, and model (2D cortical neurons vs 3D organoids at DIV20 and DIV40). Bar colors indicate treatment (DHT vs E2); shaded-experiments produced less than 10 DEGs. **(D)** Clustering of androgen and estrogen-responsive genes by dose across experiments. Line plots show standardized expression changes (z-score of log_₂_ fold change) for genes grouped into four major response patterns (top panels = androgen; bottom panels = estrogen; panel titles show number of genes per group). Each thin line is a gene; thick line and boxplots denote group median and distribution across doses. Non-monotonic responses are evident, with several gene clusters showing maximal repression or induction at lower nanomolar DHT. **(E)** Mean normalized enrichment scores (NES) from cross-experiment GSEA of DHT-induced transcriptional changes across seven independent RNA-seq datasets. Bars indicate mean NES for selected pathways (x-axis; positive = up-regulated, negative = down-regulated). Color denotes functional category (metabolism, maturation, proliferation, signaling). Circle size encodes the proportion of experiments in which the pathway was significantly enriched (FDR < 0.05). **(F)** E2 pathway enrichment analyzed as in (E).

We tested dose trends using a dose–response DESeq2 generalized linear model separately for DHT and E2. Low-count genes (total counts <10 across samples) were removed and dose values (nM) were treated as continuous variables. To improve GLM convergence and stability dose values were centered and scaled. The DESeq2 v.1.44.0(36) was used to fit the model with design = ∼dose scaled to test differential expression for each gene, but the overall DEG counts and effect sizes were not strongly dose-dependent for either hormone. DEG counts and effect sizes were not strongly dose-dependent for either hormone. The largest DHT effect occurred at 5 nM, whereas the largest E2 effect was observed at 100 nM (Figure 2D). Directionally, DHT slightly repressed gene expression relative to vehicle while the effect of E2 was more variable (Figure 2B). Non-linear dosage responses to hormone exposure are consistent with non-monotonic steroid pharmacology, where receptor/cofactor saturation, feedback, and parallel non-genomic signaling produce biphasic dose–response curves(37).

To confirm receptor availability at the treatment stage, RNAscope detected *ESR2* and *AR* transcripts in a subset of iNeurons (∼10–15% positive cells; Figure S2A) as well as ∼8% (*AR*) and ∼15% (*ESR2*) of cells in organoids. To exclude confounding from differences in cellular composition across replicates, we quantified organoid cell-type proportions (single-cell mapping) and observed minimal variability between replicates (Figures S2B, C), supporting that composition differences did not drive the bulk transcriptional results.

### Androgen response converges on neuronal maturation and cellular metabolism programs

To identify robust sex hormone-responsive programs, we compared transcriptional changes across doses, differentiation protocols, and model systems. The largest transcriptional response was in iNeurons treated with 5nM DHT in which up-regulated genes were enriched for metabolic pathways, including oxidative phosphorylation/aerobic electron transport and mitochondrial translation, whereas down-regulated genes were enriched for neuronal maturation programs such as axonogenesis, neurite outgrowth, and synapse organization. To assess whether this is a reproducible transcriptional response to DHT treatment across our experimental conditions, we performed a meta-analysis of seven independent RNA-seq experiments. Given the limited number of individually significant genes in any single experiment, we employed consistent cross-experimental gene set enrichment analysis (GSEA) to detect coordinated biological changes.

Our analysis revealed three major transcriptional signatures with high reproducibility across DHT treatment experiments (Figure 2E). First, we observed strong and consistent downregulation of neuronal differentiation and maturation pathways (significant in 6/7 experiments (85.7%)). Axon development (mean normalized enrichment score [NES] = -1.94, significant in 100% of applicable experiments) and synapse organization (mean NES = -1.80, significant in 100% of applicable experiments) showed the most robust suppression, with additional downregulation of neuron projection development (mean NES = -1.20) and neuronal differentiation regulatory programs (mean NES = -0.92). This coordinated suppression of neuronal maturation genes was further confirmed at the individual gene level, with key neuronal markers including *SYT2, NEFL, SYP, SYN1, MAP2,* and *DCX* consistently downregulated across experiments (Wilcoxon signed-rank test, p = 1.62×10⁻⁵). Second, we identified a marked upregulation of oxidative phosphorylation (mean NES = 1.83, significant in 6/7 experiments), indicating a metabolic shift toward mitochondrial respiration. Interestingly, glycolysis showed a trend toward downregulation (mean NES = -0.63), suggesting that DHT treatment promotes oxidative metabolism rather than aerobic glycolysis. This metabolic reprogramming was accompanied by modest increases in fatty acid metabolism (mean NES = 0.40). Third, we observed activation of MYC-driven transcriptional programs (mean NES = 1.31, significant in 5/7 experiments), indicating engagement of growth-promoting pathways. These gene pathways were suggestive of altered proliferation. Despite the expectation that iNeurons are largely post-mitotic, we found that at treatment iNeuron cultures retained a substantial neural progenitor/radial glia–like fraction (Nestin/NES ≈ 896 TPM; SOX2 ≈ 80 TPM; Vimentin/VIM ≈ 374 TPM) and a residual cycling population (e.g., TOP2A ≈ 36 TPM). Endothelial, pericyte, and mesenchymal markers were not detected, supporting a neural lineage identity for the NES+ cells. Acute androgen exposure may therefore be acting on a mixed culture comprising immature neurons and a residual NPC fraction. In line with this, we were unable to detect a meaningful effect on cell cycle progression machinery, perhaps reflecting the abundance of post-mitotic neurons. Unexpectedly, we found downregulation of androgen response genes (mean NES = - 0.84, significant in 3/7 experiments) which may reflect negative feedback regulation or ligand-induced androgen receptor desensitization, consistent with previously reported compensatory mechanisms in chronic androgen exposure(38),(39).

Similar to DHT, E2 treatment induced upregulation of oxidative phosphorylation (Figure 2F; mean NES = 1.01, significant in 6/7 experiments), confirming that both sex hormones promote mitochondrial metabolic activity in cultured neurons. However, the magnitude of this effect was approximately half that observed with DHT (1.01 vs 1.83), suggesting quantitative differences in metabolic reprogramming between the two hormones. In contrast to DHT’s selective metabolic activation, E2 showed modest downregulation of other metabolic pathways including glycolysis (mean NES = -0.46), fatty acid metabolism (mean NES = -0.12), and cholesterol homeostasis (mean NES = - 0.32), indicating fundamentally different metabolic profiles between the two sex hormones.

In stark contrast to DHT, E2 treatment showed strong suppression rather than activation of proliferative programs. E2F targets were markedly downregulated (mean NES = -1.81, significant in 4/7 experiments), representing the strongest transcriptional effect observed with E2 treatment aside from oxidative phosphorylation. MYC target gene sets also showed consistent suppression (MYC Hallmark set V1 mean NES = -0.91, significant in 6/7 experiments; MYC Hallmark set V2 mean NES = -0.83, significant in 3/7 experiments). This opposing effect on growth signaling pathways suggests fundamentally different cellular responses to androgens versus estrogens in our mixed neuronal cultures, with E2 promoting growth suppression appropriate for post-mitotic neurons and DHT activating proliferative programs.

E2 effects on neuronal differentiation pathways were notably more modest than DHT. While regulation of neuron differentiation showed moderate downregulation (mean NES = -0.55, significant in 2/7 experiments), and negative regulation of neuron projection development was near baseline (mean NES = -0.14, significant in 3/7 experiments), these effects were substantially weaker and less consistent than the robust neuronal suppression observed with DHT treatment (mean NES = -0.92 to -1.94). This suggests that E2 is considerably less disruptive to neuronal maturation programs than DHT.

Surprisingly, canonical estrogen response pathways showed minimal activation. Early estrogen response genes exhibited only weak upregulation (mean NES = 0.16, not significant in any experiment), while late estrogen response genes were slightly downregulated (mean NES = -0.49, not significant). This muted estrogen receptor transcriptional response may reflect differences in dosing, timing, or receptor expression levels as compared to classical estrogen-responsive tissues. Notably, E2 treatment showed strong downregulation of mTORC1 signaling (mean NES = -1.13, significant in 4/7 experiments).

Together, these findings reveal that while both DHT and E2 promote oxidative phosphorylation, they exert opposing effects on proliferative programs and differentially impact neuronal maturation pathways. E2’s transcriptional profile—moderate metabolic activation coupled with strong suppression of cell cycle machinery and minimal disruption of neuronal differentiation—may represent a more permissive environment for maintaining differentiated neuronal function as compared to DHT’s de-differentiating effects. The dramatic divergence in E2F/MYC pathway regulation (strongly activated by DHT, strongly suppressed by E2) suggests sex hormones may play opposing roles in regulating the balance between progenitor maintenance and neuronal maturation in the developing or plastic nervous system(22,27).

### *in vitro* androgen effects are consistent with the biology of the developing cortex

To relate pathway directionality to cell state, we tested whether androgen-responsive genes were enriched for progenitor versus neuronal signatures as derived from our scRNA-seq on 2D cortical cultures (methods). Up-regulated genes were significantly enriched for progenitor/radial glia/intermediate progenitor signatures, whereas down-regulated genes were enriched for maturing excitatory neuron signatures (Figure 3A). Together, these findings support a model in which acute androgen signaling shifts transcriptional programs toward proliferative/metabolic states and away from neuronal maturation. This is consistent with prior reports of androgenic regulation of growth and metabolism(40–43) and suggests a cell state–dependent repression of maturation programs in cortical lineages at these developmental stages.

**Figure 3.**
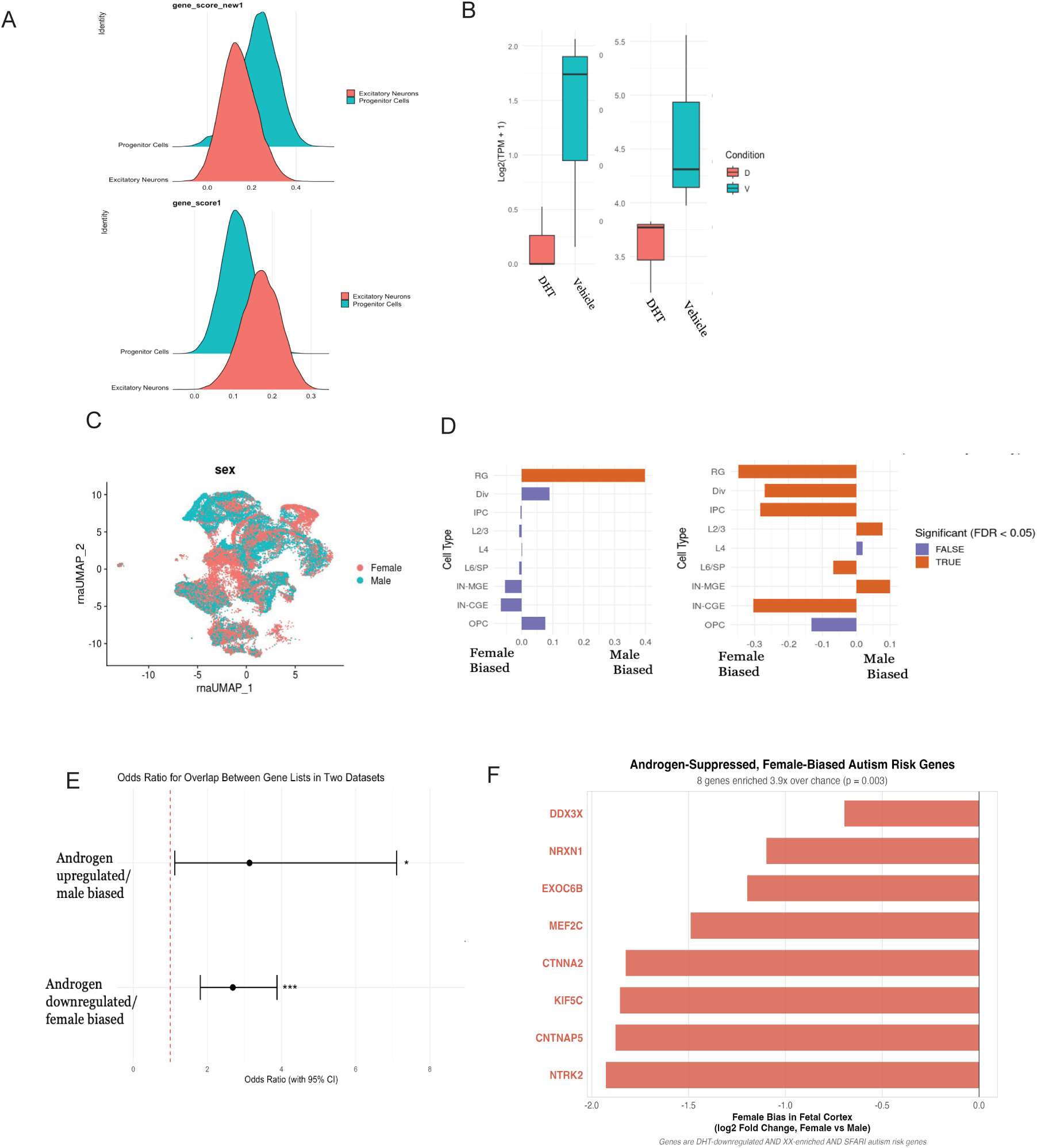
DHT transcriptional response aligns with sex-biased gene expression programs in prenatally developing cortex and is enriched amongst ASD-associated genes. (A) Distribution of DHT gene-set scores in progenitor versus excitatory neuron populations. Density plots show gene-level scores for progenitor-enriched (top) and neuron-enriched (bottom) signatures derived from prenatally developing cortex scRNA-seq. DHT-up gene scores preferentially align with progenitor signatures, whereas DHT-down gene scores align with excitatory neuron signatures, supporting a shift toward progenitor-like metabolic programs and away from neuronal maturation. **(B)** *NTRK2* is down regulated across cellular models as shown for 2D cortical neurons (left) and cortical organoids (right). **(C)** Sex differences in *NTRK2* expression across cortical cell types. Bar plot shows log_₂_ fold change (female vs male) per cell class (RG, DIV, IPC, L2/3 excitatory neurons, L4, L6/SSP, inhibitory subtypes, OPCs). Bars are colored by significance (FDR < 0.05 vs non-significant). Female-biased *NTRK2* expression is strongest in progenitor populations (RG, IPCs) and largely absent in mature excitatory neurons. **(D)** Sex differences in *AR* expression across cortical cell types. As in (C), bars denote log_₂_(female/male) expression per cell type, with significance indicated by color. *AR* is relatively enriched in radial glia and early progenitors, with minimal or no female bias in mature neurons. **(E)** Overlap of DHT-responsive genes with sex-biased genes in prenatally developing cortex. Odds ratios (with 95% CI) for overlap between DHT-up genes and male-biased genes (top) and between DHT-down genes and female-biased genes (bottom). The vertical dashed line marks OR = 1 (no enrichment). DHT-down genes are significantly enriched among female-biased genes (***P < 0.001), while DHT-up genes trend toward or show enrichment among male-biased genes. **(F)** Bar plot of the eight genes at the intersection of androgen suppression, female-biased expression, and autism risk (SFARI genes). Bars represent female bias in human prenatally developing cortex (log₂ fold change, female vs male). Genes encode proteins critical for synaptic function (*NRXN1*, *CNTNAP5, NTRK2*), axonal transport (*KIF5C*), transcriptional regulation (*MEF2C*), and RNA metabolism (*DDX3X*, X-linked).

We next sought to assess how the effect of androgen on neuronal culture models, reflects the transcriptional landscape of the developing cortex. One gene for which we observe consistent DHT down-regulation is *NTRK2*, encoding the BDNF receptor TrkB, which was downregulated by androgen in both 2D and 3D systems (Figure 3B). NTRK2 and BDNF are well-characterized targets of estrogen(17,44–46) and NTRK2 of androgen(16,47,48), suggesting that regulation of this pathway may be a key feature of sex-hormone cortical activity in general. We assessed the sex biased effect of expression of *NTRK2* in the primary human cortex (Figure 3C), within individual cell-types (Figure 3F; Wilcox-ranked sum test P<0.05). We found that *NTRK2* expression is slightly female-biased at the bulk level, with sex explaining 3.4% of expression variance in RG cells, with variation of sex-differences in individual cell-types. Notably, a significant female-bias in *NTRK2* expression is found in progenitor cell classes (RG: 34.9% higher in females; IPCs: 28.2% higher in females, both FDR < 0.001) and largely lost in mature neurons (L4 shows only 1.82% difference, non-significant; some mature neurons like L2/3 even show modest male-bias of 7.71%) (Figure 3C). This corresponds with the expression of *AR* which is largely restricted to radial glia (Figure 3C). This supports our model-system observation that *NTRK2* is down-regulated by androgen, as the expectation is that higher androgen in males would decrease *NTRK2*, leading to a female-bias in expression. Additionally, given NTRK2’s role in neuronal development, it is plausible that its downregulation would correspond with decreased maturation and increased progenitor proliferation(49–51).

We next asked if the genome-wide effects of androgen are reflected in transcriptional sex-biases in the developing cortex, reasoning that androgen responsive genes should have a directional relationship with gene expression. To this end, we used our pre-publication analysis of sex differential gene expression(29) finding that those genes which are downregulated by androgen are enriched in female-biased genes (Figure 3E; OR = 2.53, p = 5.5×10⁻⁶). Conversely, androgen upregulated genes are enriched in male-biased genes (Figure 3E; OR = 2.92, p = 0.021).

To investigate potential links between androgen-mediated sexual differentiation and neurodevelopmental disorder risk, we tested whether genes at the intersection of androgen regulation and sex-biased expression were enriched for autism risk genes. Strikingly, we found that genes both downregulated by androgen and showing female-biased expression in preantally developing cortex were significantly enriched for SFARI autism risk genes (8 of 34 genes intersecting androgen treatment and female-biased expression, 23.5%; OR = 3.90, 95% CI: 1.52-8.89, p = 0.003; Figure 3F). This enrichment was robust across multiple statistical approaches including permutation testing (p < 0.001, 1000 iterations) and remained significant across different SFARI confidence thresholds. The eight overlapping genes—*CNTNAP5, CTNNA2, DDX3X, EXOC6B, KIF5C, MEF2C, NRXN1,* and *NTRK2*—encode proteins critical for synaptic function, axonal transport, and neurodevelopmental transcriptional programs, with six classified as high-confidence SFARI genes (Scores 1/2). Notably, this enrichment was specific to the female-biased direction; androgen-upregulated, male-biased genes showed no significant enrichment for autism risk (OR = 3.21, p = 0.30). These findings suggest that disruption of androgen-suppressed, female-biased developmental programs may contribute to autism pathogenesis and could underlie the well-documented male bias in autism prevalence.

## Discussion

In this research, we provide a human-centric view of sex hormone action during cortical development, bridging data from primary human cortex and complementary culture models. We show that *ESR1/ESR2/AR* are detectable but sparse in mid-gestation cortical lineages, and that acute DHT/E2 exposures induce restricted, yet directionally consistent shifts in biochemical pathways. Androgen signaling reproducibly increases oxidative metabolism programs and suppresses axonogenesis and synapse organization, particularly in settings containing progenitor-like cells. These observations support a model in which sex hormones act selectively and, in a cell-state dependent fashion in developing cortical lineages, rather than broadly reprogramming the genome. Therefore, if sex hormones create transcriptional differences they will be in specific genes within specific cell types rather than having broad roles across the genome or across many cell types.

Our meta-analysis revealed a striking and highly reproducible suppression of neuronal maturation programs with concurrent upregulation of oxidative metabolism and MYC-regulated proliferative pathways across all DHT treatment conditions. The coordinated downregulation of axon development, synapse organization, and dendritic maturation pathways suggests an active shift towards a proliferative state rather than simply impaired maturation, as has been noted in cellular physiology.(22) This interpretation is supported by the high consistency scores (>85%) and near-universal statistical significance across independent experiments, indicating a robust biological response rather than technical variation.

The diametrically opposed transcriptional responses to DHT versus E2 treatment reveal a fundamental mechanistic divergence in how sex hormones regulate the balance between progenitor maintenance and neuronal differentiation. While both hormones converge on oxidative metabolism—albeit with differing magnitudes—their regulation of cell cycle machinery points to distinct effects on the neural progenitor populations present in our mixed cultures, particularly in cortical organoid systems. DHT’s activation of MYC and E2F targets coupled with suppression of neuronal maturation genes suggests active maintenance or expansion of the progenitor pool at the expense of neuronal differentiation. This profile is consistent with a role for androgens in preserving neural stem cell populations or blocking their differentiation. In contrast, E2’s suppression of proliferative machinery while sparing neuronal maturation programs suggests promotion of cell cycle exit and permissiveness toward differentiation—essentially the opposite developmental trajectory.

Importantly, while we detect expression of all 3 primary sex hormone receptors, to varying levels, expression of aromatase is largely undetectable. The prevailing hypothesis has long been that testosterone produced during the prenatal development of males is predominately aromatized to estrogen acting through ESR1 to affect genome activity, as is seen in mouse models, particularly subcortically. Our findings support an emerging role for testosterone via conversion to DHT acting directly on genomic targets in the developing male cortex(12,22), both in expression and localization of AR and α-reductase 1 as well as in the more robust response to DHT observed in model systems. Nonetheless, our experiments fail to address the biological context of why there is higher abundance of ESR2 than ESR1 and AR in the developing cortex. While ESR2 is classically defined as a ligand-activated nuclear receptor, evidence suggests that it can associate with chromatin and modulate transcription in a ligand-independent, context-dependent manner(52,53). Whether this occurs in this tissue requires further investigation.

Sex differences in neurodevelopmental conditions are unlikely to arise from a single mechanism. Our data support one contributory route: transient androgen signaling in progenitor-rich cortical environments biases transcriptional programs toward proliferation and metabolism while suppressing neuronal maturation. The organizational effects of perinatal androgen exposure could operate through the mechanisms we observe—prolonged progenitor maintenance, delayed or incomplete neuronal differentiation, and metabolic reprogramming that favors proliferative over mature neuronal states. Conversely, neuroprotective effects often attributed to estrogens may operate partly through promoting appropriate cell cycle exit and differentiation, preventing the accumulation of immature cells that could disrupt circuit function.

The enrichment of DHT-downregulated genes in synaptic and neurite development modules, as well as in ASD and NDD gene sets, is consistent with this model but does not imply direct causality. Rather, we propose that hormone-dependent transcriptional modulation acts alongside both autosomal and X-linked disorder gene effects, placental biology, and environmental factors to shape sex-biased developmental trajectories.

The therapeutic implications extend to regenerative medicine and neurological disorders. If androgens maintain neural progenitor populations, controlled androgen signaling might enhance endogenous neurogenesis or improve outcomes in cell replacement therapies—but only if temporally limited and followed by differentiation cues. Conversely, androgen blockade or estrogen supplementation might promote differentiation of transplanted cells or endogenous progenitors. Understanding sex hormone regulation of the progenitor-to-neuron transition could enable more sophisticated control of neural repair strategies, with sex-specific optimization of hormone-based interventions.

Key limitations include low receptor abundance, acute exposure paradigms that prioritize immediate transcriptional responses over sustained remodeling, and mixed cellular states in culture that can dilute cell-type-specific signals. Limitations in antibody quality prevented comprehensive receptor occupancy profiling, and line-specific variability (e.g., AR polymorphisms) was not explored. While scRNAseq consistency argues against composition as the primary driver, residual heterogeneity remains possible. We also did not quantify media ligand stability or intracellular hormone levels, which could contribute to non-monotonic dose effects.

Our findings replicate across 2D neurons and 3D organoids at multiple doses and developmental stages, strengthening generalizability within cortical lineages. Nonetheless, *in vitro* systems lack the full cellular complexity of the *in vivo* system, and receptor-positive cells comprise a minority population. These factors constrain effect sizes and may shift the balance of genomic versus non-genomic signaling. The modest overlap of single-gene DEGs across conditions underscores the value of pathway-level summaries for robust inference.

Together, these results support a view in which sex hormones serve as selective modulators—tuning developmental programs within the constraints of receptor availability, enhancer state, and cell identity. In the human developing cortex, androgen signaling appears to favor metabolic/progenitor programs over neuronal maturation during specific windows, providing a mechanistic foothold for understanding sex-biased developmental trajectories without invoking global genomic rewiring.

## Materials and Methods

### 2D cortical neuronal culture

#### Dual-SMAD cortical differentiation

The iPSC line M11 (male, XY; private collection, Huda Zoghbi, Baylor College of Medicine) was used for single-cell RNA-seq (scRNAseq) analysis. Karyotype and mycoplasma status were verified prior to differentiation by the Zoghbi lab and reported to be 46, XY and mycoplasma negative. Pluripotent cells were cultured on Cultrex BME matrix (R&D systems) in mTeSR Plus (STEMCELL technologies). For differentiation, cells were sub-plated in E6 medium and differentiated using dual SMAD inhibition(54) (500nM LDN-193189 2HCL, 10uM SB431542, 5uM XAV-939; Selleckchem), with NPCs sub-plated after rosette formation (day 12-19) and sub-plated in N2+B27 media (Neurobasal, GlutaMAX, L-Ascorbic acid 2-phosphate, N2 and B27) on 0.01% poly-L-ornithine (Sigma-Aldrich) + 20ug laminin (R&D systems). Neuronal maturation was induced in neuronal differentiation medium (Neurobasal, GlutaMAX, B27, 20ng/ml BDNF, 100μM L-Ascorbic acid 2-phosphate, 10ng/ml GDNF, 10μM DAPT). DAPT was withdrawn after 5 days. Cultures were validated by qPCR for cortical lineage markers (PAX6, NES, TUBB3, SLC17C1) and by scRNAseq (see “Single-cell RNA-seq”).

#### NGN2-induced iNeurons

The i3Neuron line derived from WTC-11 iPSC via NGN2 insertion at the AAVS1 locus(33) was used for direct conversion into immature glutamatergic neurons. iPSC cells were maintained as described above. For differentiation, iPSC’s were plated at 4x10^6^ cells in pre-differentiation media (DMEM/F12, NEAA, N2, 10ng/mL NT-3, 10ng/mL BDNF, 1ug/mL Laminin, 10nM Y-27632). 2μg/mL Doxycycline (DOX) was used to induce expression of NGN2. Post-induction, cells were subjected to a half-media change daily, with the addition of fresh DOX. On day 3, cells were dissociated and re-plated on poly-D-lysine (PDL)-coated plates in differentiation media (0.5X DMEM/F12, 0.5X Neurobasal-A, 1X NEAA, 0.5X GlutaMAX, 0.5X N2, 0.5X B27-VA, 10ng/mL Nt-3, 10ng/mL BDNF, 1μg/mL laminin, 2μg/mL DOX) with full media change and DOX withdrawal on day 6. iNeurons were maintained in differentiation media thereafter with weekly media changes. Culture identity was confirmed by qPCR of markers (TUBB3, SLC17C1), immunofluorescence (TUBB3), and bulk RNA-seq (see “Bulk RNA-seq”). Only male (XY) lines were used to avoid potential confounding from X-inactivation erosion in female pluripotent lines(55).

### 3D cortical organoid culture

Feeder-independent human embryonic stem cells (WA01) were obtained from the Wisconsin International Stem Cell Bank (WiCell). Cells were maintained at 37 °C, 5% CO₂ on GFR Matrigel–coated plates in StemFlex medium (Thermo Fisher Scientific) using standard feeder-free culture conditions. Cultures were passaged using 0.7 mM EDTA in Ca²⁺/Mg²⁺-free DPBS when 70–80% confluent. All lines were routinely monitored for pluripotent morphology and confirmed to be mycoplasma-free. Cerebral organoids were generated from hESCs essentially as described in(31) with minor modifications. Briefly, confluent colonies were dissociated to single cells with Accutase (Sigma-Aldrich), and embryoid bodies (EBs) were formed by seeding 9,000 cells per well in low-attachment U-bottom 96-well plates in low bFGF hES medium supplemented with Y27632 (50 µM). EBs were maintained for 3 days in hES medium without bFGF or ROCK inhibitor and then transferred individually to low-attachment 24-well plates containing Neural Induction Medium (N2-based). After 4–5 days of neural induction, aggregates displaying a radially organized, translucent neuroepithelium were embedded in Matrigel droplets (∼30 µl; Corning) on Organoid embedding sheets (stem cell technologies #08579) and cultured in Neural Induction Medium for a further 1–2 days. Two days after Matrigel embedding, organoids were transferred to 60 mm dishes and cultured in improved differentiation medium without vitamin A (IDM−A; 1:1 DMEM/F12: Neurobasal, supplemented with N2, B27–vitamin A, GlutaMAX, insulin, MEM-NEAA, 2-mercaptoethanol and penicillin–streptomycin). Where indicated, CHIR99021 (3 µM) was included for 3 days to promote early neuroepithelial expansion. Medium was then switched to CHIR-free IDM−A and changed every 3–4 days. On day 18, organoids were transferred to an orbital shaker in a 37 °C, 5% CO₂ incubator and maintained in IDM−A with medium changes every 3–4 days. Around day 20, organoids were split across two dishes to reduce density, and residual Matrigel was gently removed by mechanical disruption to improve tissue architecture. Organoids were cultured in improved differentiation medium with vitamin A (IDM+A; DMEM/F12: Neurobasal with N2, B27+vitamin A, GlutaMAX, insulin, MEM-NEAA, ascorbic acid, HEPES, penicillin–streptomycin, 2-mercaptoethanol), supplemented with dissolved Matrigel (1:50 v/v) to promote cortical plate formation. Medium was changed every 3–4 days. Between days 50 and 60, cortical plates became evident as dense, radially organized zones at the organoid surface; organoids were analyzed at the indicated time points. All media supplements (N2, B27, GlutaMAX, insulin, Antibiotic–Antimycotic) and basal media (DMEM/F12, Neurobasal, high-glucose DMEM) were from Thermo Fisher Scientific unless otherwise stated. rhFGF (R&D), Y27632 (VWR), CHIR99021 (Tocris) and ascorbic acid (Merck) were prepared as stock solutions according to manufacturers’ instructions and stored as aliquots at −20 °C.

### Hormone treatment

17β-estradiol (E2), dihydrotestosterone (DHT), and testosterone [Sigma-Aldrich] were dissolved in 100% ethanol to prepare concentrated 10mM stocks freshly on the day of exposure and equilibrated at 37°C until fully dissolved. Hormones were serially diluted in treatment media (differentiation media modified with phenol-red free Neurobasal and F-12). Exposure doses were 5, 10, 20, and 100 nM (iNeurons) and 10, 20, 100 nM (organoids) for 24 h. Vehicle controls received an identical final ethanol concentration (0.01%). In 2D cultures, hormones were diluted in media and applied via full media change. In organoids, hormone dilutions were added directly to conditioned media. After 24 h, treatment media were removed, cultures were washed twice with PBS, and cells/tissues were collected for downstream assays. All dosages were tested in 3 biological replicates, except for the organoid exposures of 10nM and 100nM.

### Single-cell RNA-seq

For 2D cultures, cells were washed twice in PBS and dissociated in Accumax (Thermo Fisher) for ∼30 min, gently lifted, and triturated to single-cell suspension. Similarly, after PBS washes, organoids were incubated briefly (∼5 min) in Accumax and dissociated by gentle trituration. In both cases, single-cell suspensions were filtered through 40 µm strainers, viability assessed, and ∼15,000 cells loaded per assay. Libraries were prepared (10x Genomics Chromium v3) and sequenced by the NYU Genome Technology Core (NovaSeq X+ 10B 100 cycle flowcell). Raw reads were processed with Cell Ranger and aligned to GRCh38. Downstream analysis was performed in Seurat V5: quality filtering (nFeature_RNA, nCount_RNA, % mitochondrial), normalization (SCTransform or log-normalize), PCA/UMAP, clustering (resolution), and annotation via known markers (RG= SOX2, HES1; Neuronal cell types= CUX1, RELN; Microglia = TMEM119)(29,32,56). Cell cycle scoring used Seurat cc.genes.updated.2019 where indicated. Label transfer or mapping to reference prenatal cortex datasets(29,56) was performed as described in Results.

### Bulk RNA-seq

Cells/tissues were lysed, and RNA was extracted using the RNeasy kit (Qiagen), quantified by Qubit, and assessed for integrity by Agilent TapeStation. All libraries had RIN 9–10. Poly(A)-selected, stranded paired-end libraries were prepared at the NYU Genome Technology Core and sequenced on a NovaSeq X (10B) using 2×100 bp reads, targeting ∼40 million paired reads per sample. FASTQ files quality were assessed by FastQC v.0.12.1 and summarized by MultiQC v. 1.25.2(57). Trimmomatic v.0.40(58) was used to remove adapter sequences and low-quality bases. The trimmed reads were aligned to GRCh38.p13(59) using STAR v.2.7.11.b(60), followed by generation of gene and transcript expression levels utilizing Salmon v. 1.10.3(61). GENCODE v38(62) annotations was used to filtered Gene and transcript count matrices to only keep protein-coding genes or transcripts. Non-coding features such as mitochondrial features, histone features, ribosomal features, pseudogenes, and non-coding RNAs (miRNA, snoRNA, and lncRNA) were excluded based on the annotation file. Afterwards, differential expression analysis was performed by DESeq2 v.1.44.0(36) with design = ∼ group and significance thresholds FDR (Benjamini–Hochberg) < 0.05. To compare DHT and estrogen transcriptional potency, we counted differentially expressed genes (padj < 0.05, Wald test) across seven independent experiments per hormone. Statistical comparisons between hormones were performed using Wilcoxon rank-sum test on experiments with detectable responses (≥1 DEG). Response rate was defined as the proportion of experiments producing ≥10 DEGs. All analyses were performed in R studio. To identify reproducible transcriptional changes across multiple DHT treatment experiments, we performed gene set enrichment analysis (GSEA) using the fgsea R package on seven independent RNA-seq datasets. For each experiment, genes were ranked by their DESeq2 Wald test statistic, and enrichment was assessed against Hallmark gene sets from MSigDB and selected Gene Ontology Biological Process terms related to neuronal function. Normalized Enrichment Scores (NES) were calculated for each pathway in each experiment, with pathways considered significantly enriched at FDR-adjusted p < 0.05. To assess cross-experiment consistency, we calculated mean NES values across all experiments and determined the proportion of experiments in which each pathway reached statistical significance. Pathways with |mean NES| > 1.0 and significance in >50% of experiments were considered robustly regulated. For gene-level consistency analysis, we evaluated directional concordance of log_2_ fold changes across experiments. Statistical significance of gene set-level directional bias was assessed using Wilcoxon signed-rank tests, with genes grouped by functional category (metabolism, proliferation, neuronal maturation). Pairwise comparisons between categories were performed using Wilcoxon rank-sum tests with Benjamini-Hochberg correction for multiple testing.

### RNAscope

Organoids were fixed in 4% paraformaldehyde for 90 min, washed twice in PBS, and embedded in paraffin using standard protocols. 2D cultures were grown on chamber slides, washed twice with PBS, and fixed in 4% paraformaldehyde for 15 min. RNAscope was performed using the ACD Bio automated protocol on the Leica BOND RX with probes against ESR2, AR, and SOX2 (Biotechne: ESR #1802921; AR #400491; SOX2 #400871). Sections (∼10 µm) were cut by the NYU Histology Core and processed immediately. Images were acquired on a Leica Polaris and processed in ImageJ. Quantification was performed as percent area positive per field or percent positive cells per cell type. Five sections/field per sample and biological replicates are reported in the figure legends.

## Primary tissue analysis

### Single-Cell RNA-Sequencing Analysis of Mid-Gestation Cortex

#### Cell-type enrichment analysis

To test whether *AR*, *ESR1*, or *ESR2* were enriched in specific cell types, we performed hypergeometric tests comparing the proportion of expressing cells (expression > 0) in each cell type versus all other cells, with Benjamini-Hochberg FDR correction for multiple comparisons across cell types in previously published Multiome data from mid-gestation cerebral cortex (GW 16–24(29)).

#### Steroidogenic enzyme expression

We extracted normalized expression values for *CYP19A1* (aromatase), *SRD5A1* (5α-reductase type 1), and *SRD5A2* (5α-reductase type 2) from the same dataset to assess local metabolic capacity for testosterone conversion. Detection rates and mean expression levels were calculated for each cell type.

#### Single-Nucleus RNA-Sequencing Analysis Across Development Dataset and preprocessing

We analyzed published snRNA-seq data from Velmeshev et al.³³ comprising 709,372 high-quality nuclei from human prefrontal cortex (BA9) spanning 40 developmental stages from prenatal development (4th gestational month, ∼16 post-conceptional weeks) through late adulthood (54 years), accessed via CELLxGENE. Expression values represent log₁₀(counts per 10,000 + 1) as provided in the original normalized data.

#### Quality control

Given the known sparse expression of nuclear hormone receptors in brain, we calculated detection rates (% nuclei with expression >0) for each receptor across all stages and compared expression to reference housekeeping genes (GAPDH, ACTB, MALAT1) to contextualize expression levels.

#### Developmental trajectory analysis

For each developmental stage, we calculated mean receptor expression across all nuclei. Stages were ordered chronologically and grouped into three periods: prenatal + 1st Year (4th gestational month through 12 months postnatal; n=14 stages, 502,533 nuclei), Childhood (2-10 years; n=6 stages, 63,049 nuclei), and Adolescence/Adult (12+ years; n=20 stages, 143,790 nuclei). Trajectories were visualized using LOESS smoothing (span = 0.75).

#### Statistical analysis

Receptor expression levels were compared using paired Wilcoxon signed-rank tests across all 40 developmental stages, with Cohen’s d calculated to assess effect sizes. Sex differences were evaluated using paired Wilcoxon tests on stage-matched male and female samples (27-30 stage-sex combinations per receptor depending on data availability). P-values were adjusted using the Benjamini-Hochberg FDR procedure to correct for multiple comparisons (three receptors tested). Statistical significance was set at FDR < 0.05.

#### Power analysis

To assess statistical power for detecting sex differences given low expression levels, we calculated detectable effect sizes for paired Wilcoxon tests with n=27-30 matched pairs at α=0.05 and power=0.8. Our analysis had 80% power to detect Cohen’s d ≥ 0.55 (medium effect), but was underpowered to detect small effects (d < 0.4). Cell-type-specific analyses with increased sample size per group would be needed to detect subtle sex differences.

All analyses were conducted in R (v4.3.0) using Seurat (v5.0.0), dplyr (v1.1.0), tidyr (v1.3.0), ggplot2 (v3.4.0), and rstatix (v0.7.2)

---

**Supplementary Figure 1.**
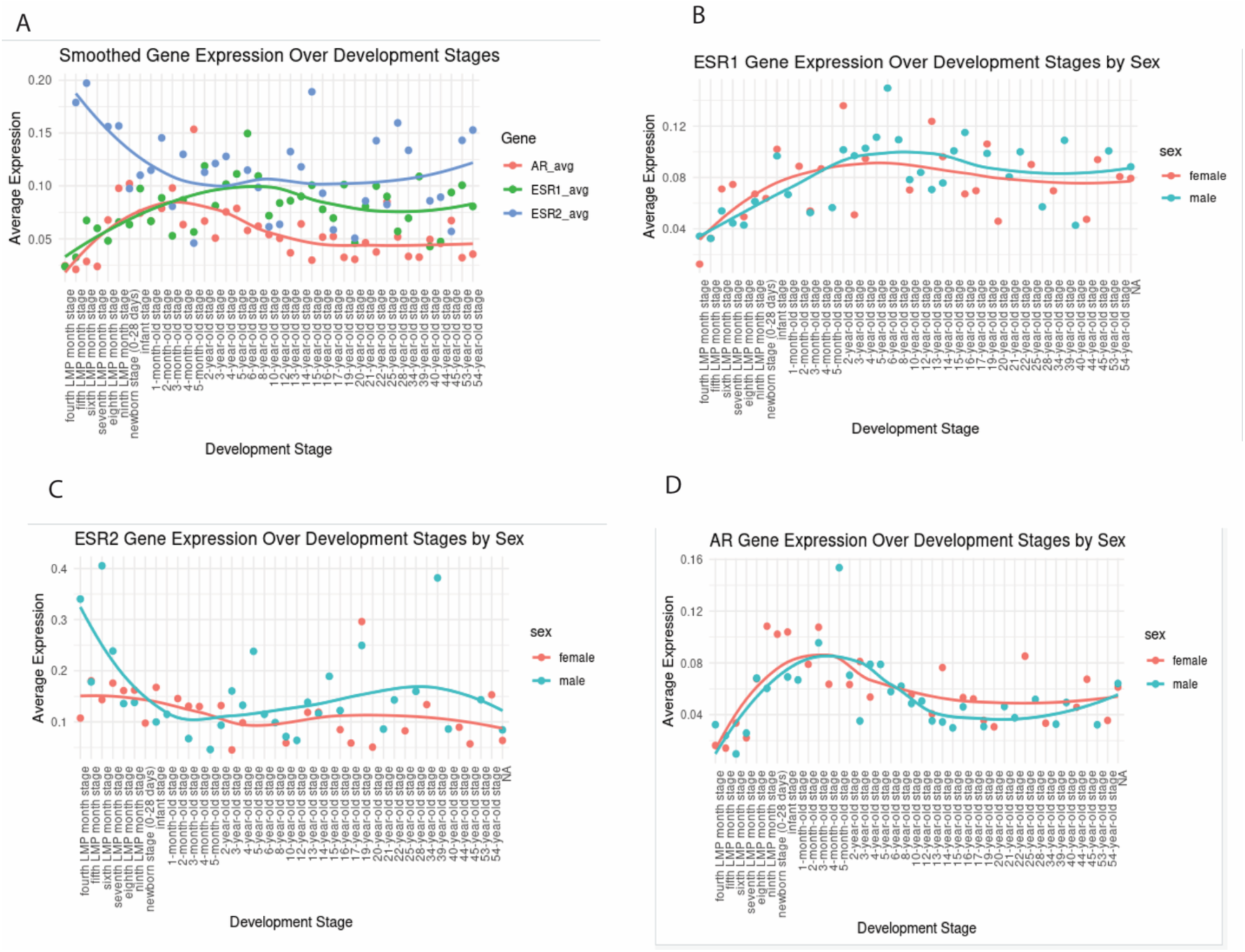
Sex hormone receptor expression dynamics across human cortical development. **(A)** Smoothed gene expression trajectories for *AR* (red), *ESR1* (green), and *ESR2* (blue) across developmental stages from early prenatal development through adulthood in primary cortex. Points represent individual samples; curves show LOESS-smoothed trends. **(B)** *ESR1* expression across development separated by biological sex (female, red; male, blue). **(C)** *ESR2* expression across development by sex. **(D)** *AR* expression across development by sex. Development stages span from early prenatal timepoints through postnatal stages. Expression values represent average normalized counts.

**Supplementary Figure 2.**
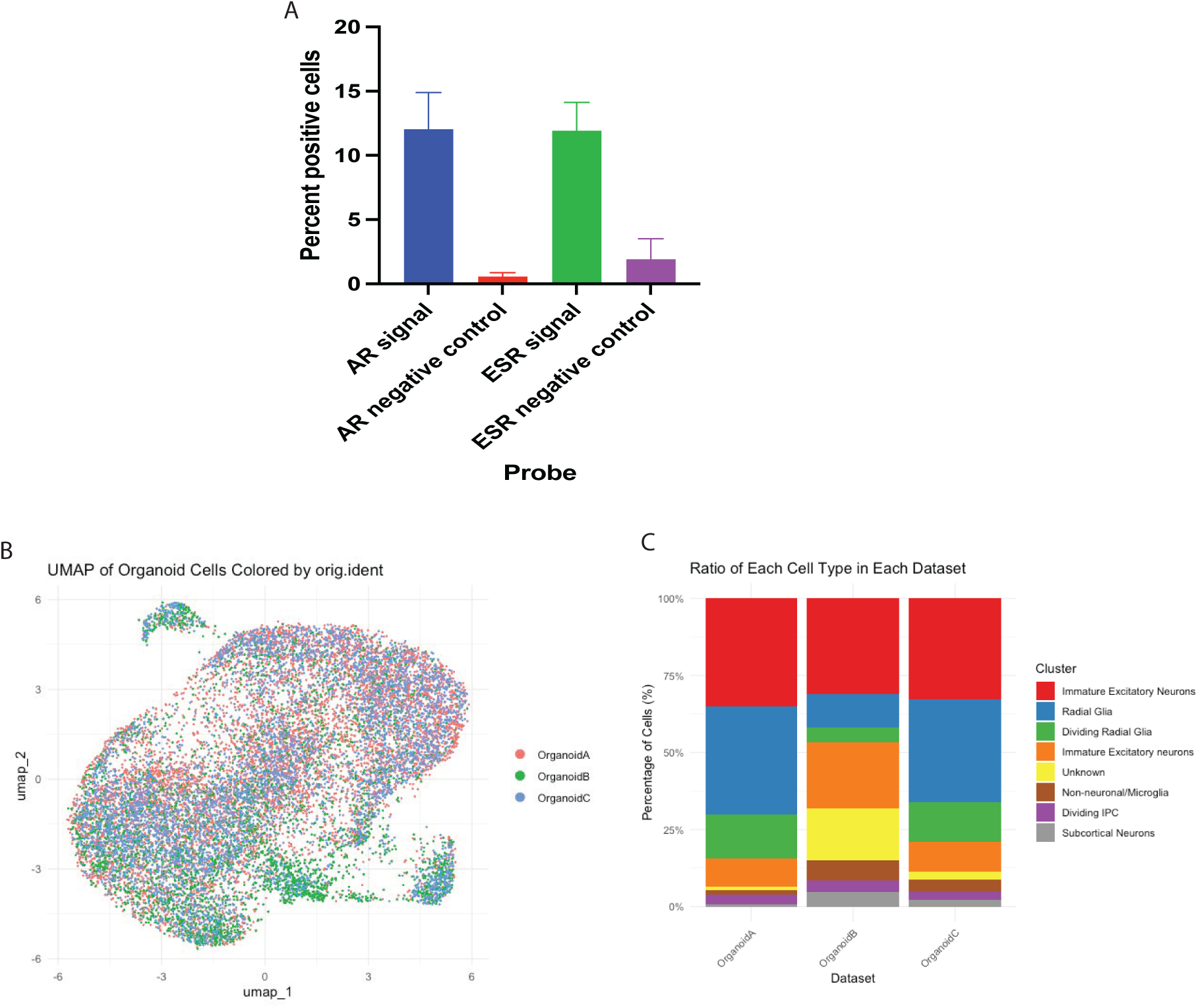
Validation of model composition for gene expression assays. **(A)** Quantification of *AR* and *ESR2* mRNA expression by RNAscope fluorescence *in situ* hybridization in iNeuron cultures. *AR* signal-positive cells: ∼12%; *ESR* signal-positive cells: ∼11%. Negative controls show minimal background (<2%). Error bars represent s.e.m. from technical replicates. **(B)** UMAP visualization of integrated single-cell RNA-seq data from three independent organoid batches (Organoid A, red; Organoid B, green; Organoid C, blue), demonstrating successful batch integration and consistent cell type identification across replicates. **(C)** Cell type composition across the three organoid batches showing reproducible proportions of major cell types as defined in figure 1.

